# tidytcells: standardizer for TCR/MHC nomenclature

**DOI:** 10.1101/2023.09.21.558778

**Authors:** Yuta Nagano, Benjamin Chain

## Abstract

T cell receptors (TCRs) underpin the diversity and specificity of T cell activity. As such, TCR repertoire data is valuable both as an adaptive immune biomarker, and as a way to identify candidate therapeutic TCRs. Analysis of TCR repertoires relies heavily on computational analysis, and therefore it is of vital importance that the data is standardized and computer-readable. However in practice, the usage of different abbreviations and non-standard nomenclature in different datasets makes this data pre-processing non-trivial. `tidytcells` is a lightweight, platform-independent Python package that provides easy-to-use standardization tools specifically designed for TCR nomenclature. The software is open-sourced under the MIT license and is available to install from the Python Package Index (PyPI).

## 1 Introduction

T cells are an important immune cell population that help orchestrate the vertebrate adaptive immune system. They express T cell receptors (TCRs) on their cell surface (fig 1a), which allows them to recognize and respond to antigens presented on the surfaces of other cells via the Major Histocompatibility Complex (MHC). Each T cell clone has a specific antigenic stimulus that it can respond to, often termed a T cell’s “cognate antigen”. The great range of target specificity is made possible by the fact that each new T cell clone generates its own unique TCR via a stochastic process of somatic gene rearrangement termed VDJ recombination.

**Figure 1:**
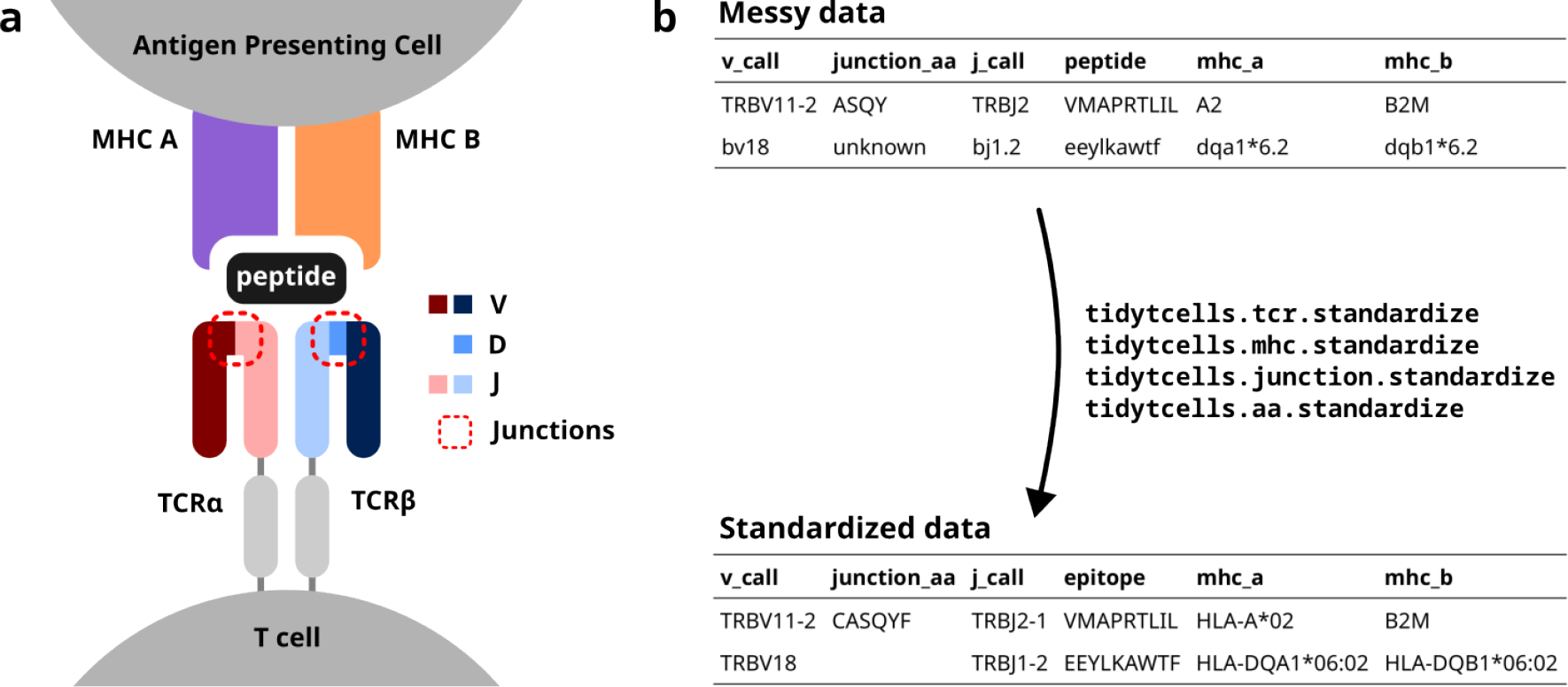
**a)** A diagram of a TCR interacting with a pMHC. The V, D and J minigenes comprising each TCR chain are shown by color. The red dotted lines point out the junction sequences of both TCR chains. **b)** An illustration of how `tidytcells` can help clean TCR data. By using `tidytcells`, non-standard nomenclature in the “messy data” is corrected, and any invalid values are filtered out.

Advances in high-throughput parallel sequencing allow large numbers of T cells isolated from blood or tissue to be sequenced. This gives us a snapshot of which T cells exist in the immune system of an individual at a given time, together with their frequency in the population, which is referred to as the individual’s TCR repertoire. Because T cell clones proliferate after recognising their cognate antigen, TCR repertoire data has proven useful as an adaptive immune biomarker in various contexts, from cancer [2, 9, 11, 13] to SARS-CoV-2 infection [3, 5, 8, 12], and there is growing excitement that TCR repertoire data can be used as a sensitive yet minimally invasive diagnostic biomarker for many other transmissible and non-transmissible diseases [14]. TCR repertoire data may also be exploited for therapeutic purposes, for example in the context of cellular therapies by contributing to the identification of TCRs with reactivities against clinically relevant targets [7, 10].

The TCR is a heterodimer, created by imprecise somatic recombination of one of a set of V and J minigenes (alpha and gamma chains), or V, J and D minigenes (beta and delta chains) (fig 1a). The current convention is to represent TCR sequence data by specifying, for each of the two chains that comprise it, which variable (V) and joining (J) genes are used, and what the aminio acid sequence is of the junction region (also known as the complementarity-determining region 3, or CDR3) between the V and J genes (fig 1b). Because of junctional imprecision, this sequence is not template driven, and cannot be aligned to the germ line sequence. In many cases, for example where populations of cells are lysed before sequencing, alpha/gamma to beta/delta chain pairing is unresolved, and indeed only one chain (typically the TCR beta) may be sequenced. In many studies, TCR sequences are further annotated by their cognate peptide/MHC (fig 1b).

Although the immunology community has generally converged to a common format of TCR data representation and has developed an international standardised nomenclature [1], TCR data in practice still contain variation due to several issues. These include (fig 1b):

- The use of non-standard TCR/MHC gene symbols
- The inclusion of non-functional TCR genes when one is only interested in data for functional TCRs
- Differing levels of TCR/MHC gene resolution-for example, some data may resolve TCR genes to the level of the allele (*TRAV1-1*01*) while others only to the level of the gene (*TRAV1-1*)

This variation is particularly problematic when trying to compile large sets of machine-readable TCRs for downstream computational analysis. For example, algorithms may not easily recognise that *TRAV1-1* and *TRAV1-1*01* are in fact the same TRAV. Similarly, *HLA-A*01* may not be understood as semantically identical to the abbreviated symbol *A1*.

`tidytcells` is a lightweight python package that addresses this issue by providing simple-to-use utilities to standardize TCR nomenclature. Its primary content is a set of functions that can convert non-standard TCR/MHC gene symbols into their international ImMunoGeneTics information system (IMGT)-standard versions [6]. Additionally, it provides simple functions to standardize junction and epitope amino acid sequences, as well as some other extra utilities.

`tidytcells` is available on the Python Package Index(PyPI) at https://pypi.org/project/tidytcells/. The source code is available at https://github.com/yutanagano/tidytcells under the MIT license. For more details such as the API reference, please see the documentation at https://tidytcells.readthedocs.io/en/latest/.

## 2 Method (Software Features)

We provide a high-level overview of `tidytcells`’ features below. More detailed instructions on usage may be found on the documentation page.

### 2.1 TCR gene symbol standardisation

`tidytcells` provides the function `tcr.standardize`, which takes as input a string representing a potentially non-standard TCR gene symbol, and outputs the corresponding IMGT-standardized symbol.

By default, if the input string cannot be resolved to a known TCR gene, the function outputs None. The function attempts to standardize to human TCR genes by default, but *Mus musculus* genes are also supported. Further options can be specified to exclude non-functional TCR genes, or limit the resolution of the symbols to the level of the gene (as opposed to allele). Below is a code block demonstrating the use of `tcr.standardize`.

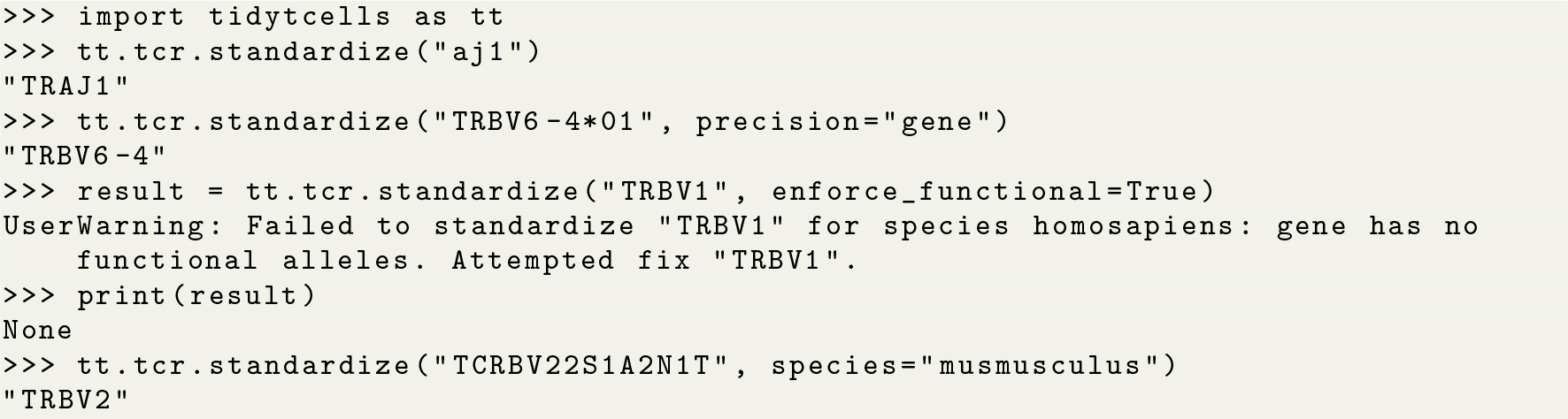

### 2.2 MHC gene symbol standardisation

A similar function `mhc.standardize` is available for standardizing MHC gene symbols. Its function signature and behaviour is essentially equivalent to its TCR counterpart.

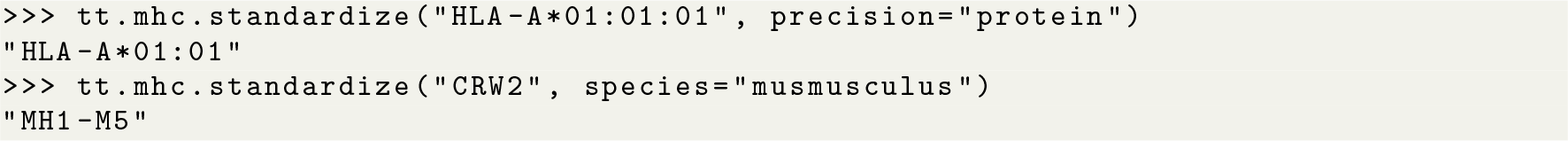

### 2.3 Junction/epitope amino acid sequence standardisation

`aa.standardize` and `junction.standardize` provide standardization utilities for amino acid sequences. aa.standardize can be used to clean generic amino acid sequence data, including epitopes, while junction.standardize provides TCR junction sequence-specific logic.

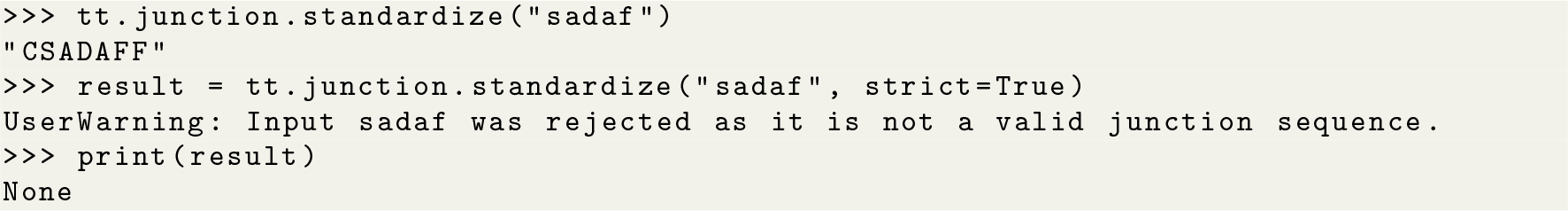

### 2.4 Extra utilities

A brief list of additional features provided by `tidytcells` is shown in table 1.

**Table 1:** A brief overview of extra utilities provided by tidytcells.

| Function | Description |
| --- | --- |
| `mhc.get_chain` | Given an MHC gene symbol, classify as alpha or beta chain |
| `mhc.get_class` | Given an MHC gene symbol, classify as class I or II |
| `mhc.query` | Query the list of all known MHC genes/alleles |
| `tcr.get_aa_sequence` | Obtain the underlying amino acid sequence of a TCR gene |
| `tcr.query` | Query the list of all known TCR genes/alleles |

## 3 Results (Application to real data)

As a test use case of `tidytcells`’ functionality, we used it in combination with the `pandas` package to clean TCR and MHC data from the Immune Epitope Database (IEDB) [4]. Where species data was available on the database, it was used. For TCR or MHC samples missing species labels, the species *Homo sapiens* was assumed. Other settings were left at default values, and no particular restrictions on gene functionality were imposed. Standardization was considered a success if the species associated with a particular value was supported, and the function managed to resolve the value to a recognised IMGT-compliant symbol.

Out of 2225 unique TCR gene symbol values found in the database, 2127 values (95.6%) were standardized successfully. Similarly, 173 of 284 MHC genes (60.9%), 301,554 of 301,670 junction sequences (99.9%) and 1825 of 1996 epitopes (91.4%) were standardized. Some examples of standardization successes and failures are shown in tables 2 and 3.

**Table 2:**
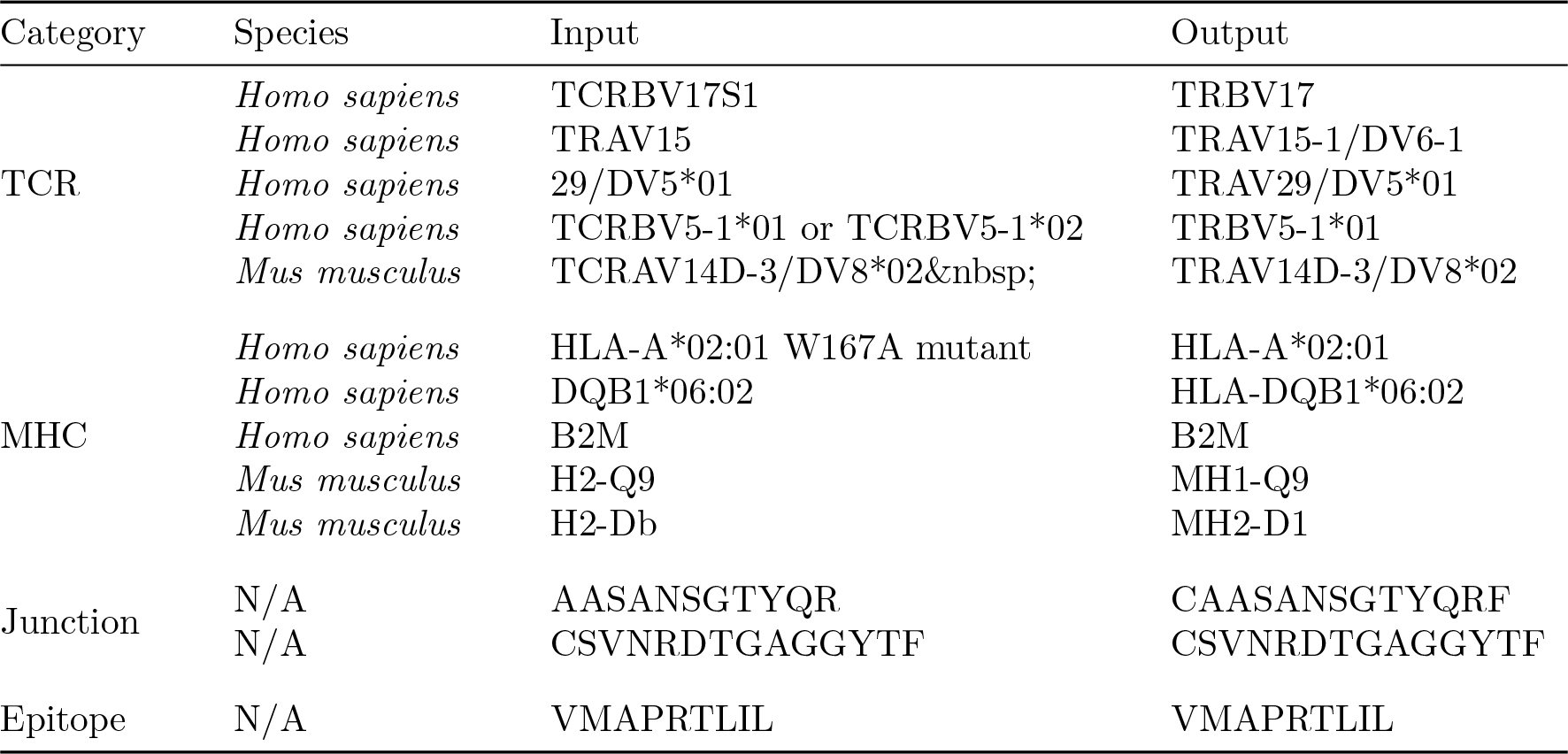
Examples of standardisation successes.

**Table 3:** Examples of standardisation failures.

| Category | Species | Input | Reason for failure |
| --- | --- | --- | --- |
| TCR | *Homo sapiens* | TCRAJ1-3 | Nonexistent gene |
|  | *Homo sapiens* | TRBV14DV4 | Nonexistent gene |
|  | *Mus musculus* | 1 | Insufficient information |
|  | *Mus musculus* | 12D-2 | Insufficient information |
|  | *Mus musculus* | TRVB13-1*02 | Nonexistent gene |
| MHC | *Homo sapiens* | HLA class II | Insufficient information |
|  | *Homo sapiens* | HLA-DQ | Insufficient information |
|  | *Homo sapiens* | human MR1 K43A mutant | Non-classical HLAs not supported |
|  | *Mus musculus* | M23I | Mutation specifier with no gene |
|  | *Mus musculus* | HLA-DRB1*04:01 | Incorrect species annotation |
| Junction | N/A | IVRVSHN*G#RDNYGQNFV | Ambiguity symbols not supported |
| Epitope | N/A | LLFGFPVYV + SCM(F5) | Peptide modification not supported |
|  | N/A | diclofenac | Not a peptide |

## 4 Discussion

As demonstrated, `tidytcells` in its current form can successfully standardise the majority of TCR/MHC data from public databases such as IEDB. However, there are still limitations to its standardization ability. Below we discuss current limitations that we as maintainers of `tidytcells` hope to address in the near future. The package is entirely open source and code contributions from the community are welcome.

Currently, when `tidytcells` encounters a string of the general form “A B” where A is a valid example of what it is attempting to standardize, it will ignore “B” and return “A” as the standardized form. This works well for cases like the first MHC success example, where the B string is a qualifier string that can be removed without fundamentally changing the underlying data. However, in cases like the fourth TCR success example where the string is of the form “A or B”, the intuitively better representation of the underlying data is to standardize to the greatest common factor of A and B (i.e. *TRB5-1* in this case). Implementing separate logic to handle these cases would improve standardization quality.

For junction sequence standardization, the default behaviour when dealing with a valid amino acid sequence that does not start with a cysteine (C) and end with a phenylalanine (F) or tryptophan (W) is to append a “C” at the beginning and an “F” at the end, and return the resulting string. The logic is implemented this way because the most common reason for these missing residues is that some data sources encode the junction as the CDR3 sequence (without the starting C and ending F/W). However, this rudimentary logic always assumes that the junction terminates with an F rather than a W. A possible improvement would be to use prior knowledge of the amino acid sequences of J genes to better predict the terminal residue. It may also be useful to provide an option to perform the reverse procedure (i.e. remove the C and F/W residues).

Other areas of potential improvement include parsing amino acid ambiguity codes and peptide modification syntax, more optional standardization constraints (e.g. specify a-priori that values should be resolved to TRAV genes/alleles as opposed to any TCR gene), support for non-classical MHC, allele imputation (if a gene has only one allele, resolve to that allele), and support for more species (only *Homo sapiens* and *Mus musculus* are currently suppoted).

## 5 Conclusion

`tidytcells` is a lightweight Python package that solves the issue of messy TCR/MHC data by providing easy-to-use utilities for standardizing TCR/MHC gene symbols, as well as general and TCR junction amino acid data. We believe this will prove to be a useful utility to the rapidly growing community of scientists who are studying the TCR repertoire.

## 6 Acknowledgments

This work was supported by the Cancer Research UK City of London Centre [grant number BCCG1C8R].

